# Expression of fibroblast activation protein-α in human deep venous thrombus

**DOI:** 10.1101/2024.02.27.582425

**Authors:** Nobuyuki Oguri, Toshihiro Gi, Eriko Nakamura, Eiji Furukoji, Hiroki Goto, Kazunari Maekawa, Atsushi B. Tsuji, Ryuichi Nishii, Murasaki Aman, Sayaka Moriguchi-Goto, Tatefumi Sakae, Minako Azuma, Atsushi Yamashita

**Affiliations:** Department of Pathology, Faculty of Medicine, University of Miyazaki, Miyazaki, Japan; Department of Radiology, Faculty of Medicine, University of Miyazaki, Miyazaki, Japan; Department of Molecular Imaging and Theranostics, Institute for Quantum Medical Science, Quantum Life and Medical Science Directorate, National Institutes for Quantum Science and Technology (QST), Chiba, Japan; Medical Imaging Engineering, Biomedical Imaging Sciences, Division of Advanced Information Health Sciences, Department of Integrated Health Sciences, Nagoya University Graduate School of Medicine, Nagoya, Japan

**Keywords:** deep vein thrombosis, fibroblast, fibroblast activation protein-α, organization

## Abstract

**Background:** Fibroblast activation protein-α (FAP), a type-II transmembrane serine protease, is expressed during wound healing, in cancer-associated fibroblasts, and in chronic fibrosing diseases. However, its expression in deep vein thrombus (DVT) remains unclear. Therefore, in this study, we investigated FAP expression and localization in DVT.

**Methods:** First, we pathologically accessed aspirated thrombi of patients with DVT (n=14), classifying thrombotic areas as follows; fresh, cellular lysis, endothelialization, fibroblastic reaction. Endothelialization and fibroblastic reaction were defined as organizing reactions. We immunohistochemically examined FAP-expressed areas and expressed cells. Second, we analyzed FAP expression in cultured dermal fibroblasts.

**Results:** All the aspirated thrombi showed a mixture of at least three of the four thrombotic areas. Specifically, 83 % of aspirated thrombi showed fresh and organizing reactions within each thrombus. Immunohistochemical expression of FAP was restricted in organizing area. Further, FAP expression in the thrombi was mainly found in vimentin-positive or α-smooth muscle actin-positive fibroblasts in double immunofluorescence. Some CD163-positive macrophages also showed FAP expression. FAP mRNA and protein levels significantly increased in cultured fibroblasts with low- proliferative activity under 0.1% fetal bovine serum (FBS) than that in fibroblasts under 10% FBS. Fibroblasts cultured in 10% FBS showed a significant decrease in FAP mRNA levels following supplementation with hemin, but not with thrombin.

**Conclusions:** The heterogeneous composition of the venous thrombi suggests the existence of a multistep thrombus formation process in human DVT. Further, fibroblasts or myofibroblasts may express FAP in organizing process in DVT. FAP expression may increase in fibroblasts with low-proliferative activity.

## Introduction

Deep vein thrombus (DVT) consists of erythrocytes, fibrin, and platelets undergoing time-dependent organizing processes, such as cellular lysis, endothelialization, macrophage infiltration, fibroblastic reactions, and replacement with collagen tissue. The analysis of the pathological characteristics of DVT has revealed the existence of a relationship between the time after the onset of thrombosis and the degree of the organizing process [1–3]. The pathological heterogeneity of thrombus contents has also been mentioned in previous reports [4, 5]. However, detailed evaluations of the pathological heterogeneity within each thrombus are limited.

Fibroblast activation protein-α (FAP), a type-II transmembrane serine protease, is expressed in activated fibroblasts, including wound-healing fibroblasts, cancer-associated fibroblasts (CAFs), and other activated fibroblasts, in various diseases [6]. Several studies have shown that FAP expression in CAFs contributes to cancer cell invasion, metastasis, and poor outcomes in various primary organs [6]. In the future, it is also expected that targeted FAP imaging will allow the detection of cancer sites [7–9]. Further, among non-oncological diseases, FAP expression has been reported in idiopathic pulmonary fibrosis, autoimmune diseases, liver cirrhosis, myocardial infarction, and atherosclerosis [6]. Sun et al. [10] reported that in mice, the inhibition of FAP expression improves cardiac function following myocardial infarction and prevents the overactivation of fibroblasts, thereby limiting scar formation. Therefore, FAP is an important target in the pathophysiology, diagnosis, and treatment of cancers and fibrotic diseases. Fibroblast and myofibroblast migration and proliferation in DVT are closely associated with extracellular matrix deposition and reduced fibrinolysis [11]. Previous studies have shown that the fibroblastic area in thrombotic tissues expresses metalloproteinase-9, osteopontin, hypoxia inducible factor 1α, and hypoxia inducible factor 2α [12, 13]. However, it remains unclear whether FAP is expressed in DVT.

Determining whether DVT shows FAP expression may have significant implications with respect to the pathophysiology, diagnosis, and treatment of DVT. Therefore, in this study, we aimed to investigate the heterogeneity of thrombus content and its association with FAP expression and localization in aspirated DVT in humans.

## Materials and methods

### Aspirated thrombi from patients with DVT

The Ethics Committee of the University of Miyazaki approved the study protocol (approval no. O-0684). Overall, 14 thrombi samples were obtained from 14 patients with DVT (eight males and six females; age range, 20–78 years; median age, 56 years; Table S1) who were diagnosed with DVT based on clinical symptoms, laboratory data, and clinical imaging findings. Proximal venous thrombi were aspirated using a guiding catheter (Guider Softip Guiding Catheter; Boston Scientific, Tokyo, Japan) installed at the popliteal vein toward the iliac vein (10 cases) or from the leg vein to the inferior vena cava (4 cases). X-ray venography revealed an extensive filling defect in the veins before thrombus aspiration, and a reduced filling defect after thrombus aspiration. After collection, all the aspirated thrombi were immediately fixed in 4% paraformaldehyde and embedded in paraffin for histological evaluation.

### Histological classification of thrombi into four components

In brief, 4-μm thick thrombus sections were stained with hematoxylin and eosin (H&E) and morphologically examined. We examined the pathological characteristics of the venous thrombi for time-dependent changes, including changes in fresh components, cell lytic changes, endothelialization, and fibroblastic reactions (Figure 1). These time-dependent changes were defined as reported in our previous study [4], i.e., fresh components were defined as areas with preserved cellular shape and structure without lytic changes or organizing reactions, cellular lytic changes were defined as the presence of leukocytes, predominantly neutrophils with the loss of cellular morphology, karyolysis, or nuclear fragmentation, endothelialization was defined as the proliferation of endothelium-like flat cells that lined the surface of the thrombi or the formation of venule-like structures [4]. Further, fibroblastic reactions were defined as the proliferation of spindle or asteroid fibroblastic cells accompanied by extracellular matrix deposition [4, 14]. Additionally, endothelialization and fibroblastic reactions were confirmed via immunohistochemistry for CD34 and α smooth-muscle actin (SMA), respectively [15].

**Figure 1.**
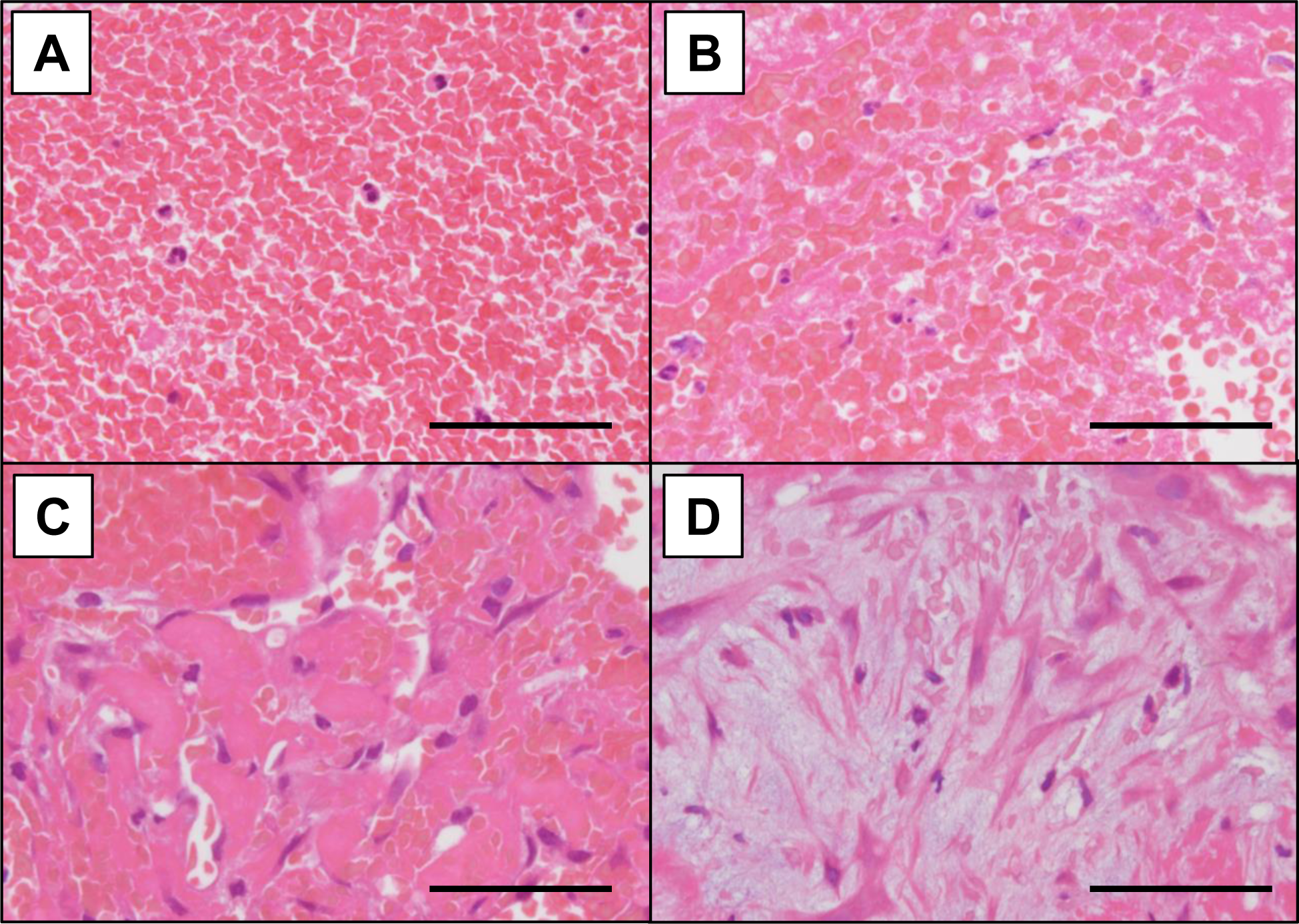
Components of human deep vein thrombus. The contents of human-aspirated venous thrombi were classified according to four histopathological types as follows: (A) Fresh thrombus, comprising erythrocytes packed within fibrin networks and inflammatory cells. (B) Cellular lysis, showing lytic changes in fibrin-surrounded leukocytes and platelet contents. (C) Endothelialization, representing embedded or lined endothelial cells in thrombus. (D) Fibroblast reactions, describing the presence of spindle or asteroid cells accompanied by extracellular matrix deposition. All the scale bars represent a length of 50 µm.

### Immunohistochemistry for human DVT

Consecutive 4-μm slices of human-aspirated thrombi were immunohistochemically stained using antibodies against glycophorin A (erythrocyte marker, mouse monoclonal antibody, clone JC159; Dako/Agilent, Santa Clara, CA, USA), fibrin (mouse monoclonal antibody, clone 59D8; EMD Millipore Corp., Burlington, MA, USA), platelet glycoprotein (GP) IIb/IIIa (platelet marker, sheep polyclonal antibody; Affinity Biologicals, Inc., Hamilton, CA, USA), CD34 (mouse monoclonal antibody, clone QBEnd10; Dako/Agilent), α-smooth muscle actin (SMA, fibroblast/myofibroblast marker, mouse monoclonal antibody, clone 1A4; Dako/Agilent), CD163 (monocyte/macrophage marker, mouse monoclonal antibody, clone 10D6, Leica Biosystems, Tokyo, Japan), FAP (sheep polyclonal antibody; Bio-Techne, Minneapolis, MN, USA), and podoplanin (mouse monoclonal antibody, clone D2-40; Dako/Agilent). Detailed information on these primary antibodies is presented in Table S2. Next, the sections were stained with EnVision anti-mouse or rabbit immunoglobulin (Dako/Agilent), or anti-sheep secondary antibodies (Jackson ImmunoResearch Inc. West Grove, PA, USA). Horseradish peroxidase activity in the thrombus sections was visualized using 3,3′-diaminobenzidine tetrahydrochloride. Thereafter, the sections were mildly counterstained with Meyer’s hematoxylin. The immunostaining controls included non-immune mouse IgG or non-immune sheep serum and not primary antibodies. Microscopic images were captured using a photosensitive color CCD camera (DS-Fi3, Nikon, Tokyo, Japan). Further, to compare the immunopositive areas between non-organizing and organizing areas, we defined organizing reactions as endothelialization and/or fibroblastic reactions. Immunopositive areas were semi-quantified using a color-imaging morphometry system (WinROOF, Mitani, Fukui, Japan) as previously described [4]. In brief, immunopositive areas were extracted as green areas using specific protocols based on color parameters in the software, namely, hue, lightness, and saturation. The data were expressed as the mean ratio of positively stained areas per thrombus area in three fields under a 20× objective lens [4]. Three fields were selected from the non-organizing and organizing areas of each thrombus.

### Immunofluorescence of venous thrombi

Immunofluorescence was performed to examine the cellular localization of FAP in representative aspirated thrombi. Briefly, the sections were stained with primary antibodies against vimentin (mouse monoclonal antibody, V9; Dako/Agilent), SMA (mouse monoclonal antibody, clone 1A4; Dako/Agilent), CD163 (mouse monoclonal antibody, clone 10D6; Leica Biosystems), and CD34 (mouse monoclonal antibody, clone QBEnd10; Dako/Agilent). Next, the sections were stained with secondary antibodies, CF488 conjugated-donkey anti-sheep IgG (Biotium, Hayward, CA, USA) and CF568 conjugated-donkey anti-mouse IgG (Biotium). The thrombus sections were then mounted using 4,6′-diamidino-2-phenylindole-containing reagents, and fluorescent images were captured using a microscope with a digital camera (DP74, Olympus, Tokyo, Japan) and merged using imaging software, Cell Sens Standard version 2.3 (Olympus).

### Fibroblast culture experiments

To examine the factors that can affect FAP expression in fibroblasts in venous thrombi, we performed fibroblast culture experiments. Specifically, primary dermal fibroblasts (normal, human, adult; PCS-201-012, ATCC, VA, USA) were cultured in Dulbecco’s Modified Eagle medium (DMEM) (Thermo Fisher Scientific Inc., Waltham, MA, USA) supplemented with 10% fetal bovine serum (FBS) (Sigma-Aldrich Co., LLC., St. Louis, MO, USA) and 1% Zell Shield (Minerva Biolabs, Berlin, Germany) (10% FBS-DMEM) at 37 °C in a humidified incubator containing 5% CO_2_.

Thereafter, we first analyzed the proliferative activity of the cultured fibroblasts. In brief, primary dermal fibroblasts (ATCC) detached via trypsinization were seeded (2 × 10^4^ cells/mL) onto an EASY GRIP Tissue Culture Dish (35 mm; Thermo Fisher Scientific) and cultured in 10%FBS-DMEM. After 24 h, the medium was replaced with 10% FBS-DMEM or DMEM (Thermo Fisher Scientific) supplemented with 0.1% FBS and 1% Zell Shield (Minerva Biolabs) (0.1% FBS-DMEM). This was followed by further incubation for 24 h, after which the cell culture supernatant was removed and the dish was fixed using methanol for 10 min at 4 °C. The dish was immunocytochemically stained using antibodies against Ki-67 (mouse monoclonal antibody, clone MIB-1, Dako/Agilent), and cell proliferation was assessed using the Ki-67 immunopositive cell ratio under a 40× objective lens.

Second, we analyzed the mRNA and protein expression levels of FAP in the cell culture. After trypsinization, the primary dermal fibroblasts (ATCC) were detached, counted, and reseeded into 6-well plates (2 × 10^4^ cells/mL) in 10% FBS-DMEM for 24 h. Thereafter, the medium was replaced with 10% FBS-DMEM or 0.1% FBS-DMEM and the cells were further incubated for another 24 h. Some of the cells incubated in the 10%-FBS-DMEM culture medium, cells were further treated with thrombin (0, 0.5, 1, 2, or 4 U/mL) (MP Biomedicals Inc., Irvine, CA, USA) or hemin (0, 10, 25, or 50 μM) for 24 h (Porcine Hemin, FUJIFILM Wako Pure Chemical Corp., Osaka, Japan). FAP mRNA and protein expression levels were assayed using qPCR and enzyme-linked immunosorbent assay (ELISA), as described in the sections below.

### RNA isolation and qRT-PCR analysis of cultured fibroblasts

Human dermal fibroblasts were cultured as already described in the previous section. Thereafter, the cell culture supernatant was removed and the cells were washed twice with 2 mL of cold PBS. Next, TRIzol reagent (Invitrogen, Carlsbad, CA, USA) was added, and total RNA was then extracted using PureLink RNA Mini Kits (Thermo Fisher Scientific). The amount of RNA was quantified using a NanoDrop 1000 spectrophotometer (ND-1000; Thermo Scientific, Rockford, IL, USA). Then to synthesize cDNA, we used the PrimeScript RT Reagent Kit (Perfect Real Time; Takara Bio Inc., Shiga, Japan). Finally, FAP and β-actin mRNA expression were determined using TB Green Premix Taq (Takara Bio). The primers used for β-actin were as follows: 5′-TGGCACCCAGCACAATGAA-3′ (forward) and 5′-TAAGTCATAGTCCGCCTAG-AAGCA-3′ (reverse). Those used for FAP were 5′-TGTATCGAAAGCTGGGTGTTTATGA-3′ (forward) and 5′-GATGCAAGGGCCAGTGATGA-3′ (reverse). Real-time PCR was performed using a LightCycler 96 instrument (Roche Diagnostics GmbH, Mannheim, Germany). FAP mRNA expression level was normalized to that of β-actin.

### Protein expression analysis using ELISA

Total protein was extracted from cultured cells using Pierce RIPA Buffer (Thermo Fisher Scientific) containing 1% Halt Protease Inhibitor Cocktail (Thermo Fisher Scientific). The protein concentration was determined using the Pierce BCA Protein Assay Kit (Thermo Fisher Scientific). Thereafter FAP protein expression level was quantified using a Quantikine Colorimetric Sandwich ELISA kit (R Systems, Minneapolis, MN, USA) and normalized to the total protein concentration.

### Statistical analyses

All data were expressed as medians and interquartile ranges. Differences between or within groups were evaluated by performing the Mann–Whitney U-test and the Kruskal–Wallis test with Dunn’s multiple comparison using GraphPad Prism software version 9 (GraphPad Software Inc., San Diego, CA, USA). Statistical significance was set at P [0.05 (n indicates the number of samples).

## Results

### Heterogeneous histological findings for human-aspirated DVT

In this study, we examined the time-dependent changes in the pathological characteristics of DVT by investigating in fresh components, cell lytic changes, endothelialization, and fibroblastic reactions (Figure 1). Table 1 shows the changes in the components of the human-aspirated DVT with time after the onset of thrombosis. From this table, it is evident that all the 14 DVT examined showed heterogeneity in thrombotic content, each comprising more than three components. Specifically, 64% of the DVTs showed all four components, 29% showed fresh, lytic, and endothelialization components, and 7% showed lytic, endothelialization, and fibroblastic reactions. Figure S1 shows the representative microphotographs of the DVT containing the fresh, lytic, endothelialized, and fibroblastic reaction components. Further, eight of the nine thrombi (89%) that initially showed fresh components still displayed this characteristic even 10 days after onset. Only the aspirated thrombus in case 2, which was aspirated 60 days after onset, showed no fresh components.

**Table 1.** Time after onset and components of human deep vein thrombus.

|  | (day) | Component |  |  |  |
| --- | --- | --- | --- | --- | --- |
|  |  | Non-organizing area |  | Organizing reaction |  |
|  |  | Fresh | Lytic | Endothelialization | Fibroblastic reaction |
| Case 1 | 5 | + | + | + | - |
| Case 2 | 60 | - | + | + | + |
| Case 3 | 10 | + | + | + | + |
| Case 4 | 9 | + | + | + | - |
| Case 5 | 14 | + | + | + | + |
| Case 6 | 30 | + | + | + | + |
| Case 7 | 11 | + | + | + | + |
| Case 8 | 8 | + | + | + | + |
| Case 9 | 30 | + | + | + | + |
| Case 10 | 6 | + | + | + | + |
| Case 11 | 7 | + | + | + | + |
| Case 12 | 14 | + | + | + | + |
| Case 13 | 26 | + | + | + | - |
| Case 14 | 30 | + | + | + | - |

### Localization of FAP in human DVT

We subjected the aspirated thrombi to immunohistochemical analysis and thereafter, compared the immunopositive areas for each component between non-organizing and organizing areas. Figure 2A shows the representative histological and immunohistochemical images of the non-organizing and organizing areas of the thrombi. The non-organizing areas predominantly consisted of erythrocytes, fibrin, and platelets. CD34, SMA, or FAP expression was not observed in the non-organizing areas. Conversely, the organizing area showed CD34-positive endothelialization as well as SMA-positive fibroblastic reactions accompanied by the infiltration of CD163-positive monocytes/macrophages. Further, FAP expression was observed in the spindle-shaped cells in the organizing areas. However, none of the DVT specimens showed podoplanin-immunopositive cells (data not shown). Further, immunopositive areas for erythrocytes, fibrin, and platelets were significantly higher in the non-organizing areas than in the organizing areas (Figure 2B), while immunopositive areas for CD34, SMA, CD163, and FAP were significantly higher in organizing areas than in non-organizing areas (Figure 2B).

**Figure 2.**
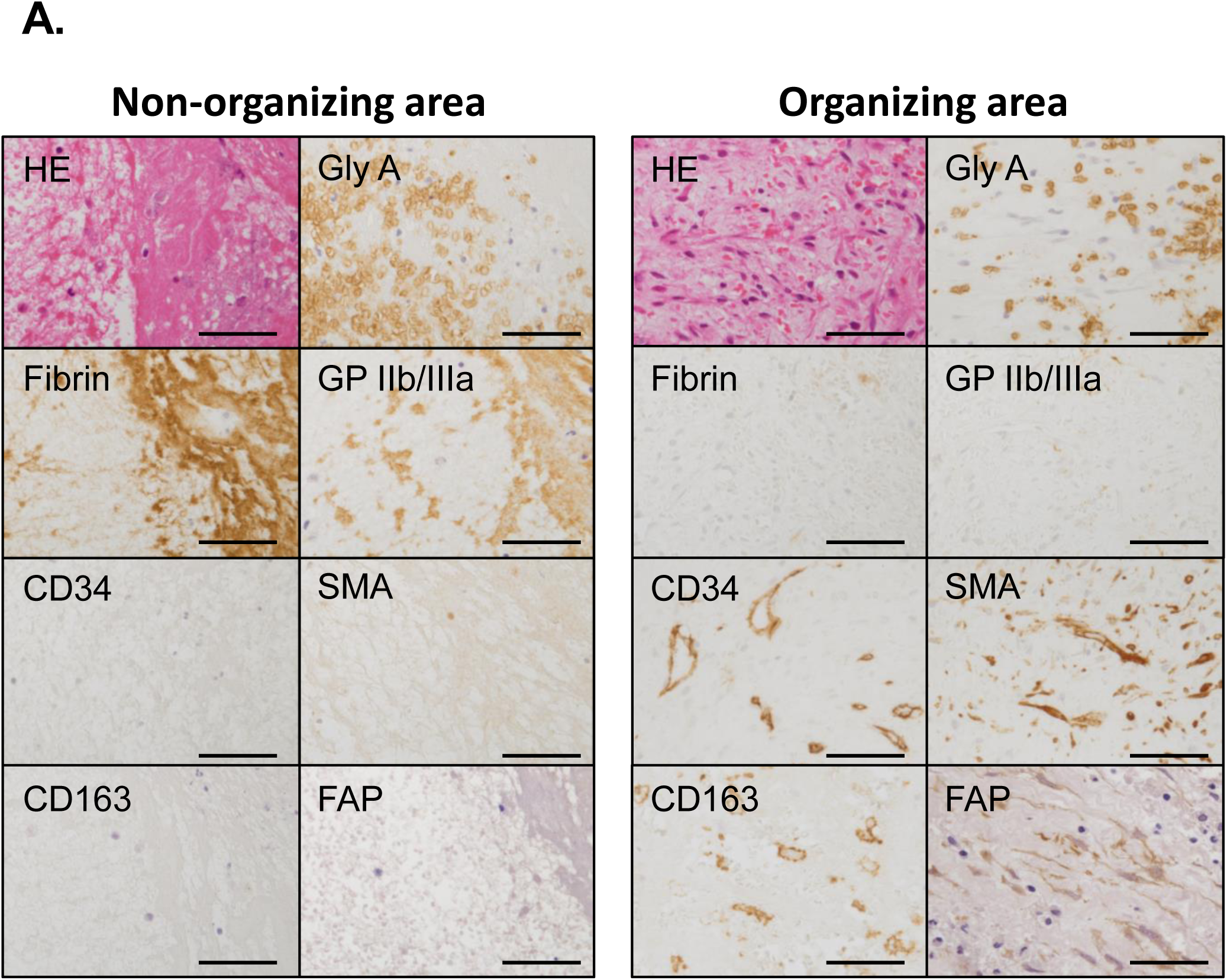
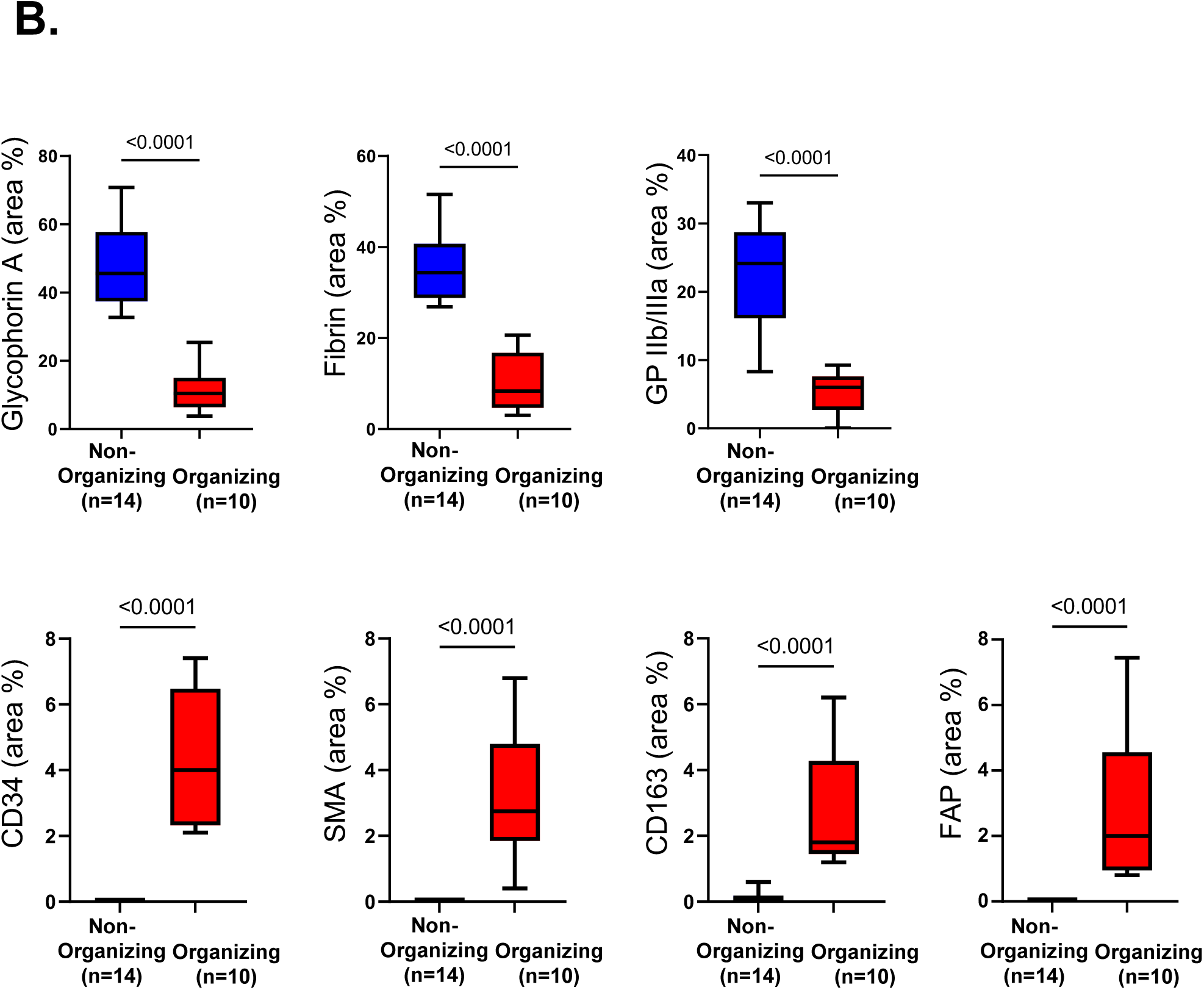
Fibroblast activation protein-α expression in human-aspirated deep vein thrombus. (A) Representative microphotographs of thrombus contents in non-organizing and organizing areas. The non-organizing area predominantly consisted of glycophorin A (Gly A, erythrocyte), fibrin, and glycoprotein (GP) IIb/IIIa (platelet). No immunoreaction was observed for CD34, α-smooth muscle actin (SMA), CD163, and fibroblast activation protein-α (FAP). The organizing area showed less immunoreaction for fibrin, GP IIb/IIIa, and glycophorin A, but showed the presence of CD34-, SMA-, and CD163-immunopositive cells. FAP-immunopositive spindle cells were also observed in the organizing area. All the scale bars represent a length of 50 µm. HE, hematoxylin and eosin stain. (B) Immunopositive area for thrombus contents in non-organizing and organizing areas. Organizing areas were defined as areas with the presence of SMA-immunopositive cells. The ratio of the immunopositive area was calculated based on the median values of the three representative fields for each thrombus under a 40× objective lens. The Mann– Whitney U test was used for statistical analysis.‏

Immunofluorescence was performed to examine the cellular localization of FAP in representative DVT specimens. The results thus obtained indicated that FAP expression predominantly occurred in vimentin- or SMA-immunopositive spindle cells, which are considered to be fibroblasts or myofibroblasts. A few CD163-immunopositive cells also showed FAP expression (Figure 3).

**Figure 3.**
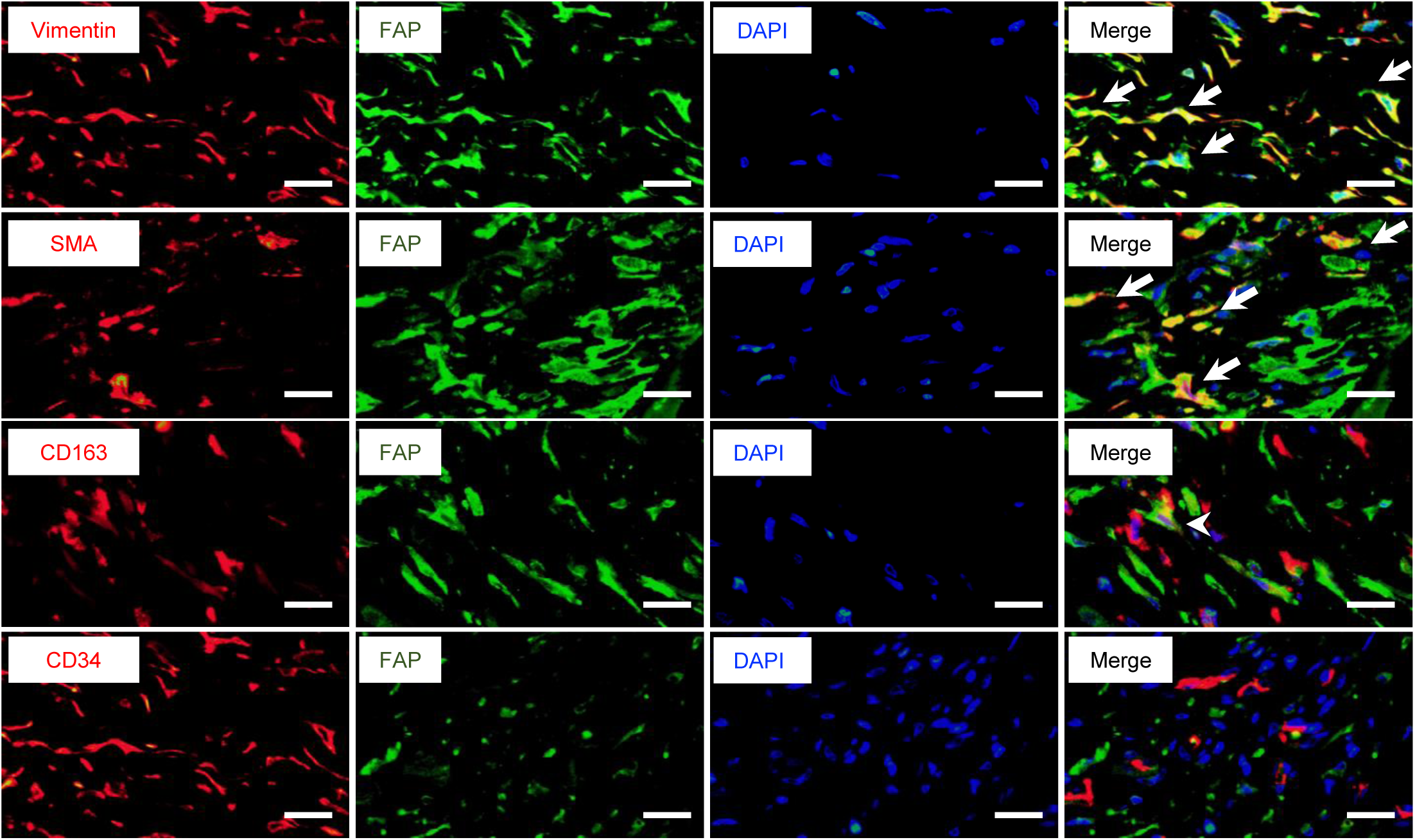
Cellular localization of fibroblast activation protein-α in aspirated human deep vein thrombus. Representative immunofluorescent images of fibroblast activation protein-α (FAP) in human-aspirated venous thrombi. The organizing areas show co-localization of vimentin and FAP in spindle-shaped cells (arrows). Smooth muscle actin (SMA)-positive cells were frequently positive for FAP expression (arrows). A few CD163-positive cells showed FAP expression (arrowhead). CD34-positive cells did not show FAP expression. All the scale bars are 20 µm in length. DAPI, 4’,6-diamidino-2-phenylindole.

### Fibroblast proliferative activity affected FAP expression

We analyzed the proliferative activity of cultured human fibroblasts supplemented with 10% or 0.1% FBS. Thus, we observed that the fibroblasts cultured in 0.1% FBS showed a significantly lower proportion of Ki-67-immunopositive cells than those cultured in 10% FBS (median value, 21% vs. 77%; Figure 4A and 4B). Further, the FAP mRNA and protein expression levels of fibroblasts cultured in 0.1% FBS were significantly higher than those of fibroblasts cultured in 10% FBS (Figure 4C and 4D). These observations indicated that fibroblast proliferative activity affected FAP expression.

**Figure 4.**
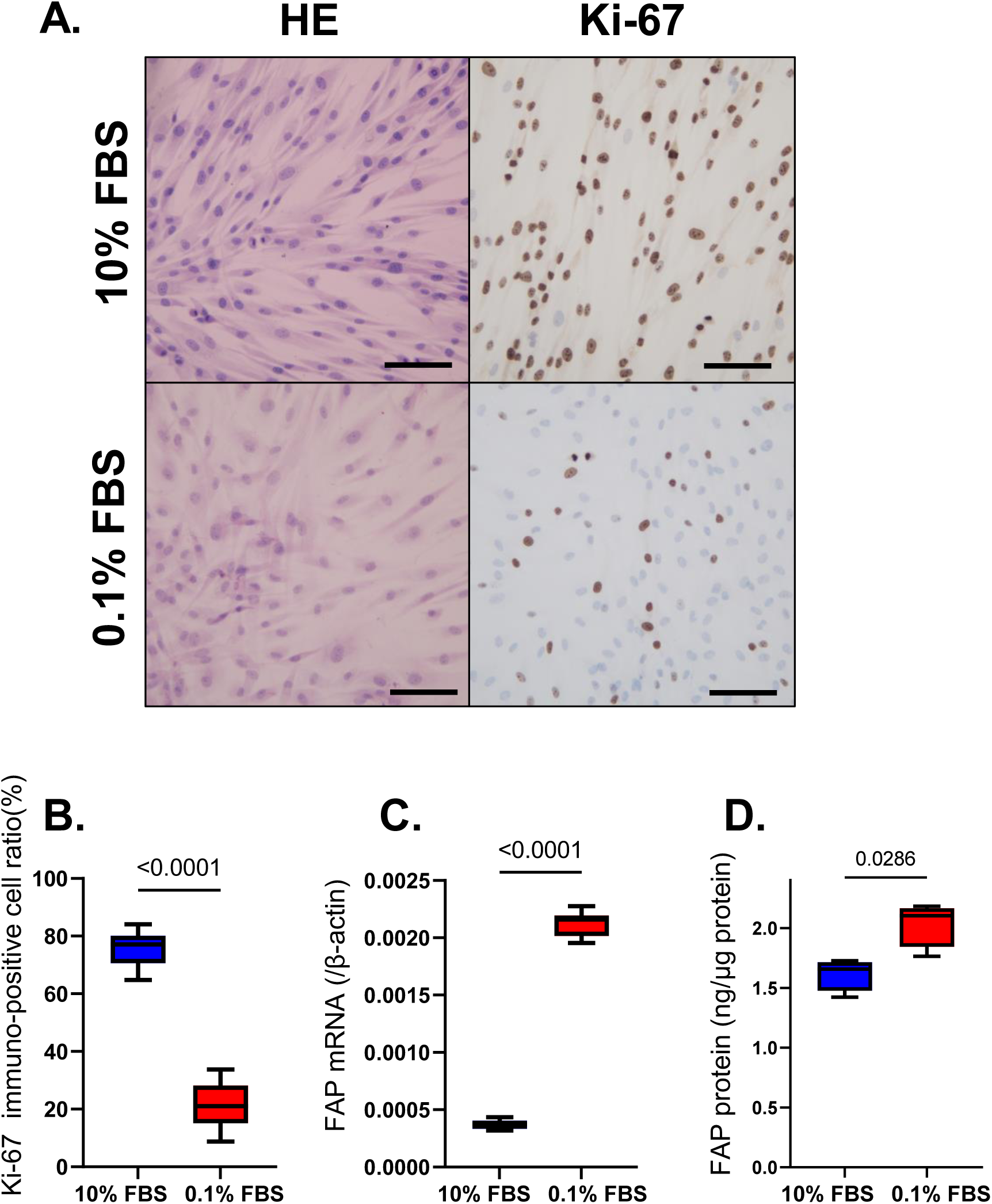
Effects of cell proliferation on fibroblast activation protein-α expression in cultured fibroblasts. Representative microphotographs of hematoxylin and eosin stain (HE) and Ki-67 immunocytochemistry (A). All the scale bars indicate a length of 50 µm. Fibroblasts cultured in 0.1% FBS for 24 h showed less-Ki-67 immunopositivity than those cultured in 10% FBS (A, B). Fibroblast activation protein-α (FAP) mRNA (C) and protein (D) levels after culturing in 10% FBS and 0.1% FBS for 24 h. Statistical analysis was realized by performing the Mann–Whitney U test (B, C, and D).

### Hemin, not thrombin affected FAP expression in cultured fibroblasts

To examine whether thrombotic factors affected FAP expression in fibroblasts, we measured FAP mRNA levels in cultured fibroblasts supplemented with different concentrations of thrombin or hemin, an iron-containing porphyrin derived from hemoglobin. The results obtained revealed no significant differences in FAP mRNA levels among the different thrombin concentrations (0, 0.5, 1, 2, and 4 U/mL; Figure 5A). Further, we observed that hemin supplementation significantly downregulated FAP mRNA expression levels in the fibroblasts (Figure 5B).

**Figure 5.**
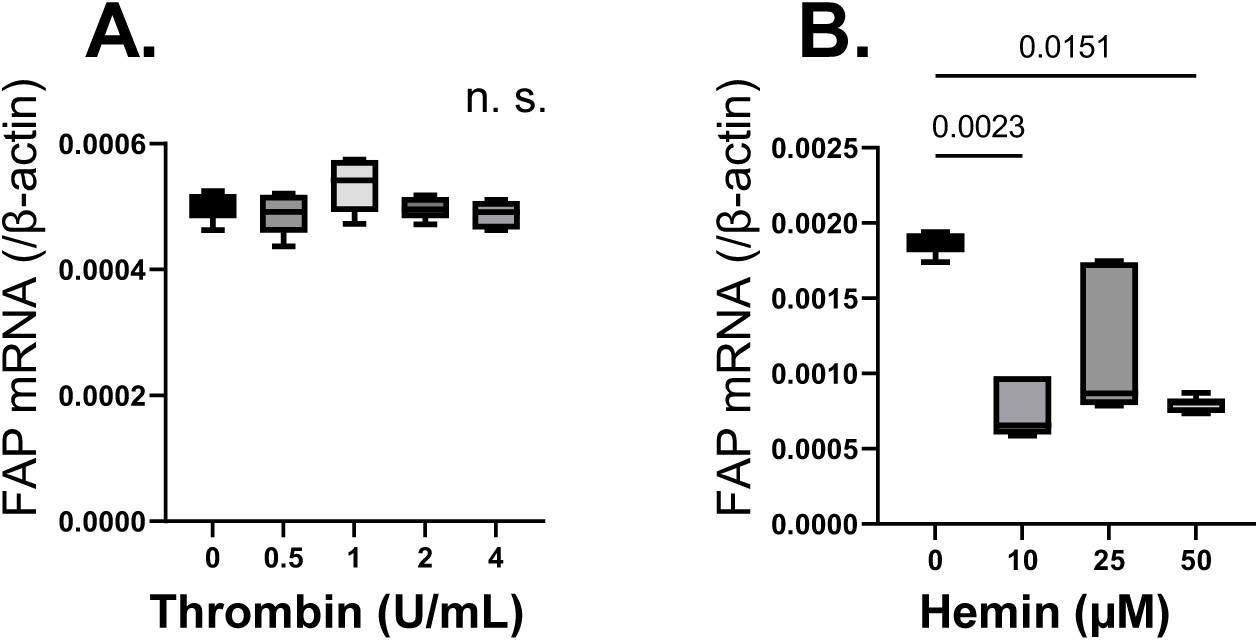
Effects of thrombin or hemin on fibroblast activation protein-α expression in cultured fibroblasts. Fibroblast activation protein-α (FAP) mRNA levels following: (A) Thrombin and (B) Hemin exposure for 24 h. The Kruskal-Wallis test with Dunn’s multiple comparison was performed for statistical analysis.

## Discussion

The results of this study showed that human-aspirated DVT is characterized by various phases of time-dependent changes and that FAP is expressed in fibroblasts/myofibroblasts in the organizing areas of these DVT. Additionally, fibroblasts with low proliferative activity showed upregulated FAP expression, while hemin supplementation resulted in a decrease in FAP expression.

DVT comprises erythrocytes characterized by large amounts of fibrin and relatively few platelets [4, 16]. An immunohistochemical study also showed that platelets and von Willebrand factor are present in DVT and pulmonary embolism (PE) [17]. In the present study, all aspirated DVT lesions showed various phases of time-dependent changes, in addition to fresh components. In previous pathological studies on human DVT, researchers evaluated the presence of citrullinated histone H3, a marker of neutrophil extracellular trap formation, in non-organizing or organizing areas or in fresh, cell lytic, and organizing components within DVT or PE [4, 18]. Their findings highlighted differences in the thrombus formation phase or cellular/organizing reactions within each DVT or PE. Furthermore, an autopsy-based study showed that cell lytic changes occur 3–8 days after DVT formation [19]. In this currentstudy, the presence of fresh components in DVT at > 10 days after onset suggests additional thrombus formation (thrombus growth) after the initial onset. These findings suggest the existence of a multistep growth phase of venous thrombus formation.

FAP expression has been reported in CAFs and activated fibroblasts in various organ diseases [6]. It has also been observed that FAP can also be expressed in non-fibroblast cells, such as endothelial cells and macrophages [6, 20, 21]. FAP expression in endothelial cells is associated with capillary-type tubule formation in vitro [20]. A subpopulation of tumor-associated macrophages was found to show FAP expression and exhibit immunosuppressive function in a mouse tumor model [21]. In the present study, we observed that in human DVT, FAP is mainly expressed in the organizing area of fibroblasts or myofibroblasts. Even though some CD163-positive macrophages showed FAP expression in the organizing area, no FAP-positive endothelial cells were observed in DVT. Podoplanin is a well-known CAF marker similar to FAP [22]. Using a mouse model, Shirai et al. [23] reported that extracellular vesicles derived from CAFs promote venous thrombus formation via podoplanin-dependent platelet aggregation. In the present study, podoplanin-positive cells were not observed in the aspirated thrombi. Therefore, FAP-positive fibroblasts or myofibroblasts in human DVT may be different from CAFs in terms of podoplanin expression, and may not promote platelet aggregation or thrombus formation via podoplanin-dependent pathway.

Factors capable of regulating FAP expression have also been reported. Brokopp et al. [24] reported that macrophage-derived tumor necrosis factor α can induce FAP in cultured human aortic smooth muscle cells. Chen et al. [25] also reported that tumor growth factor-β can induce FAP in cultured mouse fibroblasts. However, it is unclear

whether the intrathrombotic environment affects FAP expression. Interestingly, we observed that proliferative activity affected FAP expression in fibroblasts. The organizing reaction in DVT is similar to the process of wound healing, in which fibroblasts migrate and proliferate to synthesize and deposit extracellular matrix proteins. Under these conditions, cell migration and proliferation are mutually exclusive [26]. Additionally, sensing an increasing gradient of platelet-derived growth factors may switch fibroblasts from a migrating to a proliferating phenotype [27]. Pohl et al. [28] reported that thrombin and fibrin can induce fibroblast proliferation in vitro. In the present study, thrombin supplementation did not affect FAP expression in cultured fibroblasts. Therefore, FAP may be abundantly expressed in the migratory phenotype of fibroblasts in organizing thrombi. Further, hemin, the oxidized prosthetic moiety of hemoglobin, is released into the blood of patients with severe hemolytic condition [29]. Some in-vitro studies have suggested that hemin induces pulmonary endothelial dysfunction [30, 31]. We assumed the presence of hemin in DVT tissue, because we previously reported a hemolytic morphology of erythrocytes in the human DVT [1]. In the present study, we observed that hemin administration decreased FAP expression in cultured fibroblasts. Therefore, it is possible that hemin derived from degraded erythrocytes downregulates FAP expression in DVT.

FAP exhibits enzymatic activity with respect to the cleavage of α2 anti-plasmin, leading to its conversion into a more potent inhibitor of plasmin and a decrease in the rate of lysis of the fibrin clot [6, 32]. The suppression of fibrinolysis by α2 anti-plasmin contributes to stasis-induced venous thrombus formation in mice [33]. Therefore, FAP may enhance fibrin formation in DVT. However, we did not observe FAP in the fresh fibrin areas in the present study. The findings suggests that the conversion of α2 anti-plasmin rarely occurs in the fresh components of human DVT. In contrast, almost all the thrombi examined in this study showed the coexistence of fresh and organizing areas. Therefore, FAP-expressing fibroblasts/myofibroblasts may negatively regulate fibrin degradation via the conversion of α2 anti-plasmin.

^68^Ga-FAP inhibitor-46 is a radiolabeled probe for the in vivo detection of FAP [7, 8, 34], and its positron emission tomography can be used to detect DVT but not pulmonary emboli in patients with venous thromboembolism [35]. Besides, our previous studies showed that DVT and pulmonary thromboembolism are composed of heterogeneous components of fresh and organizing thrombi [2, 4]. The discordant uptake of DVT and pulmonary emboli may be due to differences in thrombus components. Further, DVT may have an organizing component; however, fresh components can detach and embolize pulmonary arteries. ^68^Ga-FAP inhibitor-46 positron emission tomography may be suitable for detecting organizing thrombi.

This study had several limitations. First, the sample size, i.e., the number of human-aspirated DVT examined, was small. Second, we confirmed the presence of FAP immunoexpression; however, the role of FAP in human DVT still unclear. Therefore, further studies are required to determine the role of FAP in thrombus organization.

## Conclusions

Histopathological composition of venous thrombi suggested the existence of a multistep thrombus formation process in human DVT. Further, we primarily observed FAP expression in fibroblasts/myofibroblasts, accompanied by the organizing process of DVT. Our results also indicated that FAP expression may be increased in fibroblasts with low proliferative activity.

## CRediT authorship contribution statement

Nobuyuki Oguri: Writing – original draft, Investigation, Visualization, Methodology, Data curation, Formal analysis. Toshihiro Gi: Writing – review & editing, Investigation, Visualization, Methodology, Data curation, Formal analysis. Eriko Nakamura: Investigation, Data curation. Eiji Furukoji: Methodology, Data curation, Supervision. Hiroki Goto: Investigation, Visualization, Data curation, Formal analysis. Kazunari Maekawa: Investigation, Deta curation. Atsushi B. Tsuji: Conceptualization, Methodology, Supervision. Ryuichi Nishii: Conceptualization, Methodology, Supervision. Murasaki Aman: Supervision, Methodology. Sayaka Moriguchi-Goto: Supervision, Methodology. Tatefumi Sakae: Supervision. Minako Azuma: Project administration, Supervision. Atsushi Yamashita: Supervision, Project administration, Conceptualization, Methodology, Writing – review & editing.

## Abbreviations list

CAF: cancer-associated fibroblast
DMEM: Dulbecco’s Modified Eagle medium
DVT: deep vein thrombosis
FAP: fibroblast activation protein-α
FBS: fetal bovine serum
GP IIb/IIIa: glycoprotein IIb/IIIa
SMA: smooth muscle actin

## Data availability

The data presented in this study are available on request from the corresponding author.

## Acknowledgments

We thank Nahoko Udatsu and Kyoko Ohashi for their technical assistance. We would like to thank *Editage (*www.editage.com) for English language editing.

## Supplemental Materials

Table S1, S2

Figure S1

