## Supplementary material for "Expression of fibroblast activation protein-α in human deep venous thrombus": Figure S1

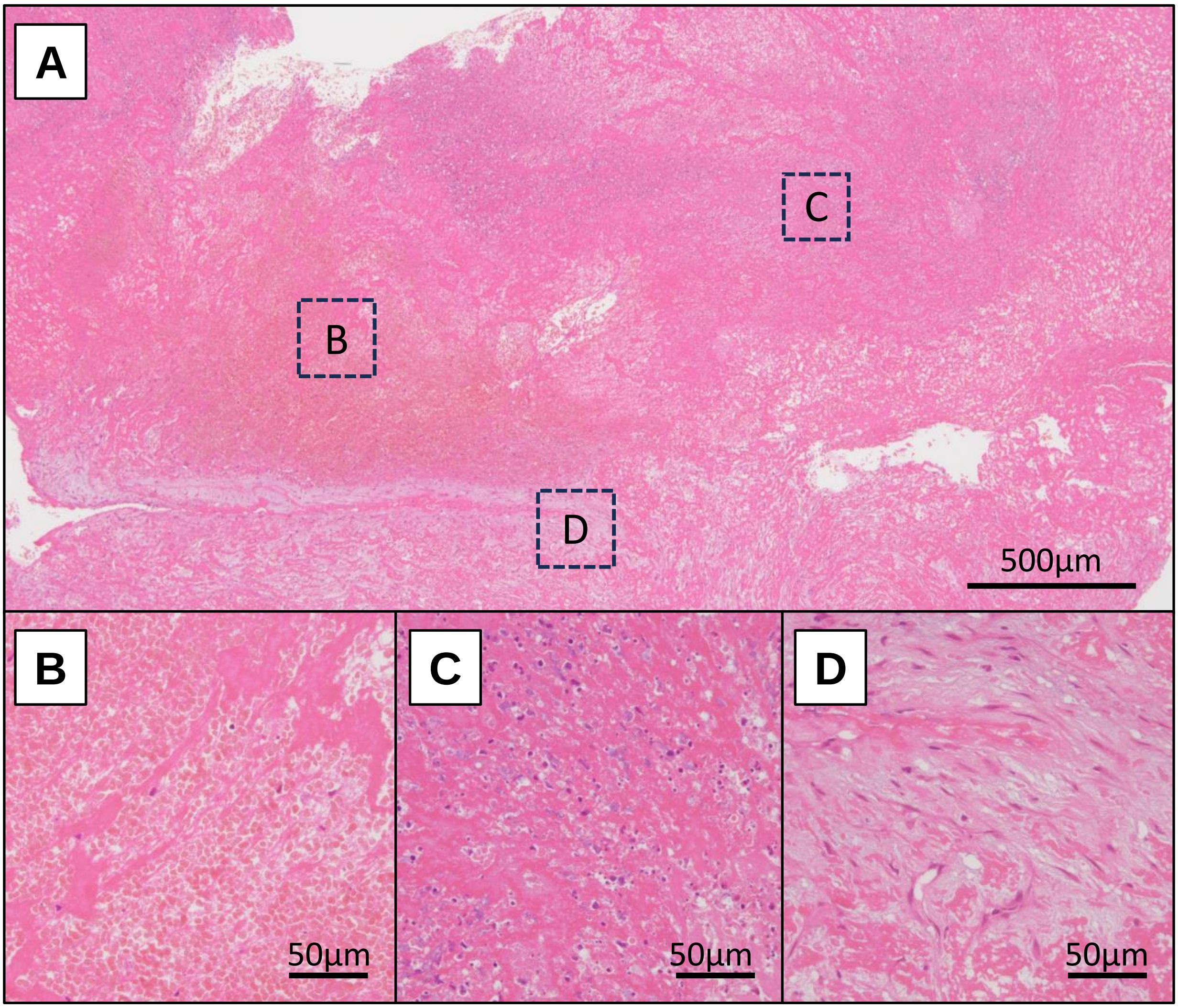


**Figure S1. Pathological heterogeneity within a deep vein thrombus.**

Representative microphotographs of heterogeneity of an aspirated venous thrombus. Mixture of fresh and organizing contents is observed within the thrombus (A, low-magnification), such as erythrocyte-rich area (B), cellular lysis of leukocytes (C), and endothelialization and fibroblastic reaction (D). High-power fields of B-D are corresponding to the insets of A.
