## Supplementary material for "Expression of fibroblast activation protein-α in human deep venous thrombus": Table S1, S2

| **Table S1. Clinical background of the patients with DVT (n=14)** | | |
| --- | --- | --- |
| Age, median (range; years) | 56 | (20-78) |
| Male sex, n (%) | 8 | (57) |
| Obesity; BMI >25, n (%) | 3 | (21) |
| Smoking, n (%) | 8 | (57) |
| Complication |  |  |
| Post-traumatic, n (%) | 4 | (29) |
| Cancer, n (%) | 2 | (14) |
| Pulmonary embolism | 6 | (43) |

BMI, body mass index

| **Table S2. Primary antibodies for immunohistochemistry/ immunocytochemistry** | | | | | | | |
| --- | --- | --- | --- | --- | --- | --- | --- |
| Antibody | Antigen/Marker | Species | Clone | Concentration | HIER | Company | Catalog number |
| Glycophorin A | Erythrocyte | mouse | JC159 | post-dilution | none | Dako/Agilent | M0819 |
| Glycoprotein IIb/IIIa | Platelet | sheep | Polyclonal | 20 μg/mL | none | Affinity Biologicals Inc. | SA2B3A-IG |
| Fibrin | Fibrin | mouse | 59D8 | 0.5 μg/mL | MW | EMD Millipore Corp. | MABS2155-25-UG |
| CD34 | Endothelium | mouse | QBEnd-10 | post-dilution | MW | Dako/Agilent | M7165 |
| α-Smooth muscle actin | fibroblast/myofibroblast | mouse | 1A4 | post-dilution | MW | Dako/Agilent | M0851 |
| CD163 | Monocyte/macrophage | mouse | 10D6 | 0.5 µg/mL | MW | Leica Biosystems | NCL-L-CD163 |
| Fibroblast activation protein α | Serin protease | sheep | Polyclonal | 20 µg/mL | none | Biotechne | AF3715 |
| Podoplanin | Podoplanin | mouse | D2-40 | 0.37 μg/mL | MW | Dako/Agilent | M3619 |
| Vimentin | Stromal cell | mouse | V9 | post-dilution | MW | Dako/Agilent | M0725 |
| Ki-67 | Cell proliferation | mouse | Mib 1 | post-dilution | MW | Dako/Agilent | N1633 |
| HIER, heat-induced epitope retrieval methods; MW, microwave | | | | | | | |
